# Long-lived mammals contain more phosphorylation sites in the SIRT6 C-terminus that enhance PARP1 interaction and resistance to oxidative stress

**DOI:** 10.64898/2026.04.22.718469

**Authors:** Jonathan Gigas, Michael E. Meadow, Jing Guo, Catherine Lan, Eric Hillpot, John C. Martinez, Gregory Tombline, Philip Bellomio, Valeria Rivera Almodovar, Kevin A. Welle, Kyle Swovick, Jennifer R. Hryhorenko, Julia Ablaeva, Sina Ghaemmaghami, Andrei Seluanov, Vera Gorbunova

## Abstract

Sirtuin 6 (SIRT6) is a protein deacetylase and ribosyltransferase that is a vital hub for maintaining epigenetic homeostasis, regulating the transcriptome, and repairing DNA double stranded breaks (DSBs). Comprehensive proteomic profiling of the SIRT6 post-translational landscape, however, remains elusive. The SIRT6 C-terminal domain contains multiple phosphorylation sites. We find that the presence and use of these sites is strongly correlated with maximum lifespan across mammals. Subsequent biochemical and *in silico* analyses revealed that SIRT6 hyperphosphorylation enhances its interaction with PARP1. Mutating the T294 phosphorylation site in human fibroblasts led to decreased survival after oxidative stress in the phospho-null T294A and improved oxidative stress resistance in the phospho-mimetic T294E. Together, these results suggest SIRT6 C-terminal phosphorylation increases stress resistance through interaction with PARP1, and that this effect has been enhanced by evolution in long-lived species.

## INTRODUCTION

Sirtuins are a family of deacetylases and mono(ADP-ribosyltransferases) that are essential for healthy aging. Broadly, it is thought that sirtuins help fend off many human age-related hallmarks like genomic instability [1, 2], telomere dysfunction [3–5], and chronic inflammation [6, 7]. SIRT6 is one member of the sirtuin family that has a particularly strong link to longevity. SIRT6 activity has been shown to be correlated with lifespan in mammals and *Drosophila* [8, 9]. For example, male mice overexpressing SIRT6 have a 30% extension in lifespan compared to wildtype mice without any observable deleterious phenotype [10, 11]. SIRT6 specifically has been implicated in many cellular processes, including DNA repair [2, 12], pluripotency [13, 14], metabolism [15], cancer [16, 17], immunity [18, 19], and gene expression [20–22]. How SIRT6 manages to be involved in so many disparate processes throughout the cell, however, is not fully understood.

SIRT6 is an early responder to sites of DNA damage, where it cooperates with PARP1 to promote repair [2]. Multiple studies have used laser micro irradiation to show that the kinetics of SIRT6 recruitment to DNA damage sites is very rapid and promotes recruitment of PARP1 [2, 12, 23]. This has established SIRT6 as an upstream regulator of DNA repair, even before the selection of a repair pathway. The mechanism of this recruitment is unclear, though it has been shown that the C-terminus of SIRT6 has a high affinity for naked DNA in addition to its core domain’s affinity for nucleosomes [24]. SIRT6 ADP-ribosylation of PARP1 enhances PARP1 activation and subsequent repair of double-stranded breaks [2, 23]. Through this mechanism, overexpression of SIRT6 stimulates both NHEJ and HR by more than three-fold. This activity is further enhanced by JNK-mediated phosphorylation of SIRT6 at serine 10 (S10) during oxidative stress, priming the nucleus for repair of oxidative DNA damage [23]. The NHEJ-stimulation capacity of SIRT6 across rodent species was shown to correlate with species maximum lifespan, suggesting that this mechanism is selected for during the evolution of extended lifespans [8].

An understudied aspect of SIRT6 regulation is its numerous post-translational modifications (PTMs). Data from a number of global proteomic discovery projects, as aggregated in the CST PhosphoSitePlus database, report that there may be 27 or more modifiable residues throughout the protein [25]. Some of these modifications have been shown to impact the function of SIRT6. For example, deacetylation of K33 by SIRT1 is important for proper SIRT6 polymerization and recognition of double stranded breaks [26]. Five phosphorylation sites were rigorously validated by *Miteva et al.* in 2014 [27], and these remain the most commonly annotated PTMs for SIRT6 in the database. Only two of these have published functions: phosphorylation at S10 by JNK leads to active participation in DSB repair [23], and AKT-mediated phosphorylation of S338 is reported to lead to MDM2-mediated degradation of SIRT6 [28]. However, there is no unified consensus on the complete profile of SIRT6 post-translational regulation. Most of the putative SIRT6 modifications have not been validated by low-throughput experiments, and their functional significance remains unknown. For example, putative SIRT6 phosphosites at T294, S303, T305, S326, S330, and T337 are largely uncharacterized [29]. Interestingly, the proline-rich and unstructured C-terminus appears to be a hub for modifications, including four of the five phosphorylation sites reported by *Miteva et al.* [27].

In this study, we investigated the C-terminal phosphorylation of SIRT6 across mammalian species with diverse lifespans. We show that SIRT6 is highly phosphorylated in human cultured cells, that SIRT6 C-terminal phosphorylation is strongly correlated with species maximum lifespan, and that C-terminal phosphorylation enhances the interaction of SIRT6 with PARP1 thereby conferring enhanced oxidative stress resistance.

## RESULTS

### SIRT6 is heavily phosphorylated in human cells

SIRT6 consists of a short unstructured N-terminus, globular Rossman fold domain, and long unstructured C-terminus as visualized in the predicted AlphaFold2 structure (Figure 1A). Multiple phosphorylation sites had been detected in the SIRT6 C-terminus in proteomics studies (Figure 1B) [25, 27]. Interestingly, many of these sites contain proline in the -1 or +1 positions (Figure 1C) which may suggest kinase specificity. Throughout the literature, Western blot of SIRT6 appears as a doublet with one band at 42 kDa (slightly higher than the predicted 39 kDa) and another slower mobility band at 46 kDa. Here, we confirm that SIRT6 migrates as two distinct bands across a variety of human cell lines (Figure 1D).

**Figure 1.**
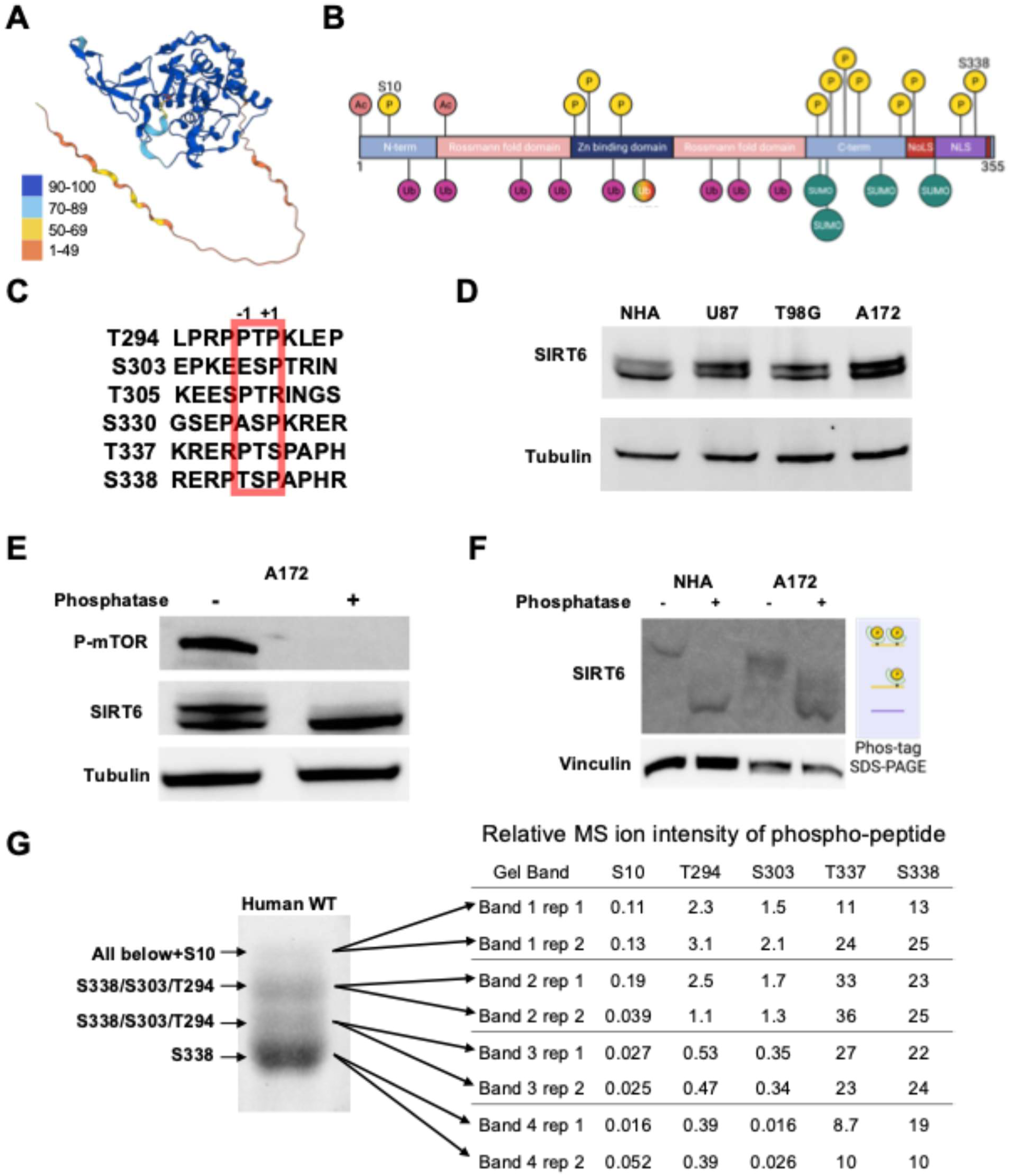
SIRT6 is hyperphosphorylated in human cells. **A)** AlphaFold structure prediction of human SIRT6 (Q8N6T7). Structure is colored according to the pLDDT scale of prediction confidence, included as a reference. **B)** Combined map of identified SIRT6 PTMs in the literature and through global proteomic discovery projects. Acetylations – S2, K33. Phosphorylations – S10, T146, Y148, T162, T294, S303, T305, S310, S326, S330, T337, S338. Ubiquitinations – K17, K33, K81, K128, K160, K230, K245, K267. ncUbiquitination – K170. SUMOylations – K296, K300, K316, K332. NLS – nuclear localization sequence. NoLS – nucleolar localization sequence. **C)** Aligned human SIRT6 amino acid sequences from the eight most consistently identified SIRT6 phosphosites. The +1 position is proline on each of the sites. **D)** Western blots on cell lysates from human astrocytes (Lonza NHA) or glioblastoma (U87, T8G, A172) cell lines. **E)** Western blot on A172 cell lysates prepared with or without lambda phosphatase treatment. **F)** Western blot on cell lysates after performing SDS-PAGE in a FUJIFILM Phos-gel that dramatically slows electromobility of phosphorylated proteins[30]. **G)** Coomassie stained gel bands after extended SDS-PAGE on SIRT6 purified from HEK293T cells. Phosphorylations enriched in each band as identified by MS are labeled for reference.

We found that treating cell lysates with lambda phosphatase eliminates the separation of these bands, suggesting that differential phosphorylation of SIRT6 proteoforms is responsible for their separation (Figures 1E and S1A). This was confirmed by using a Phos-Tag gel which magnifies the electromobility shift of phosphorylated proteins [30], and revealed that all SIRT6 in cell lysates is significantly slowed in these gel conditions (Figures 1F and S1B). These results suggest that phosphorylation of SIRT6 may be ubiquitous in human cells, though this modification does not appear to cluster by cellular compartment (Figure S1C). To measure this more directly, we purified human SIRT6 recombinantly from HEK293 cells. Performing LC-MS/MS on this recombinant protein without any further enrichment for phospho-peptides revealed a significant abundance of C-terminal phosphorylations (Table 1). We then separated SIRT6 protein on a 30 cm polyacrylamide gel into four distinct and apparently differentially modified proteoforms and performed in-gel digest and LC-MS/MS on each band (Figure 1G). We observed a general increase in relative phosphopeptide abundance in bands with slower electromobility, which is consistent with their separation being due to phosphorylations (Table 2). We associated each band with a distinct set of phosphorylations enriched in it based on these data. In total, 4 proteoforms were separated, which identified differential N- and C-terminal phosphorylations and showed that SIRT6 is ubiquitously phosphorylated at S338, frequently at S303 and T294, and rarely at S10.

**Table 1.** Example MS phosphoproteomic data for recombinant SIRT6. Mass spectrometry quantitation results from a typical run of purified WT human SIRT6 produced recombinantly in HEK293T cells. Each modified vs unmodified peptide is identical with the exception of the phosphorylation, including cysteine alkylation. S326 is included as an example of the many reported phosphosites that are not usually detected in our data. The other sites in this table are the only phosphorylated peptides consistently quantified in our experiments.

| Site | MS2 Spectral Count |  | MS1 Ion Intensity |  |
| --- | --- | --- | --- | --- |
|  | Unmodified | Modified | Unmodified | Unmodified |
| S10 | 56 | 16 | 1.42E9 | 4.28E7 |
| T294 | 87 | 90 | 1.81E10 | 1.60E10 |
| S303 | 13 | 64 | 5.55E7 | 6.47E8 |
| S326 | 29 | 0 | 1.13E9 | 0 |
| S330 | 29 | 35 | 1.13E9 | 1.73E9 |
| T337 | 19 | 139 | 1.03E8 | 8.47E9 |
| S338 | 19 | 36 | 1.03E8 | 8.59E9 |

**Table 2.** Relative phosphorylation in separated SIRT6 gel bands. Relative phosphorylation for each gel band after LC-MS/MS, as calculated by dividing the MS1 ion intensity of the phosphopeptide by that of the same unmodified peptide. #1 and #2 refer two each of two replicate experiments. A general trend of increasing phosphorylation towards the slowest mobility bands is observed.

| Gel Band | S10 | T294 | S303 | T337 | S338 |
| --- | --- | --- | --- | --- | --- |
| Top #1 | 0.11 | 2.3 | 1.5 | 11 | 13 |
| Top #2 | 0.13 | 3.1 | 2.1 | 24 | 25 |
| High #1 | 0.19 | 2.5 | 1.7 | 33 | 23 |
| High #2 | 0.039 | 1.1 | 1.3 | 36 | 25 |
| Low #1 | 0.027 | 0.53 | 0.35 | 27 | 22 |
| Low #2 | 0.025 | 0.47 | 0.34 | 23 | 24 |
| Bottom #1 | 0.016 | 0.39 | 0.016 | 8.7 | 19 |
| Bottom #2 | 0.052 | 0.39 | 0.026 | 10 | 10 |

**Table 3.** Measurement of SIRT6 C-terminal phosphorylation by HIPK. Relative measurement of SIRT6 C-terminal phosphorylation was calculated by dividing the ion intensity of the phosphopeptide by the ion intensity of the unmodified peptide. WT and harmine samples were purified recombinantly from HEK293T cells with or without treatment with harmine during the production phase. The HIPK sample was purified recombinantly from E. coli and phosphorylated with recombinant HIPK2 kinase. * = the unmodified peptide was undetectable, which in all cases was accompanied with very high phosphopeptide abundance.

| Site | Treatment |  |  |
| --- | --- | --- | --- |
|  | WT | Harmine | HIPK |
| T294 | 1.12 | 0.313 | * |
| S303 | 17.6 | 3.80 | 0.0132 |
| S330 | 3.40 | 2.48 | 0.154 |
| S338 | 126 | 20.7 | * |

According to these data, SIRT6, as measured by either MS, Coomassie staining, and Western blot intensity, appears to harbor multiple C-terminal phosphorylations. Thus, we find that SIRT6 is constitutively phosphorylated on its C-terminus in human cell culture and frequently has more than one such modification.

### SIRT6 phosphorylation correlates with maximum lifespan

After identifying multiple phosphorylation sites of the C-terminus of human SIRT6, we next sought to understand if this feature is conserved across mammalian species. SIRT6 activity in DNA repair is well correlated with maximum lifespan in rodents [8], and the C-terminus of SIRT6 has significant sequence diversity throughout mammals. We downloaded SIRT6 sequences from NCBI for all mammalian species that are present in the AnAge database [31, 32]. These sequences were aligned to confirm the appropriate positioning of the C-terminus relative to each other (Figure 2A), the results of which show excellent sequence conservation up to position 298, after which there is a large amount of variation in both sequence and length. We then counted the number of predicted phosphorylation sites according to the [ST]P motif. The sites were all present on the C-terminus, except for the near-invariant S10 site. We observed a significant positive correlation between the relative number of putative C-terminal phosphorylation sites and species maximum lifespan (Figure 2B). This correlation is robust to phylogenetic correction using a phylogenetic least squares regression, implying that the relationship between predicted sites and lifespan is independent of the evolutionary relationship of the species to each other.

**Figure 2.**
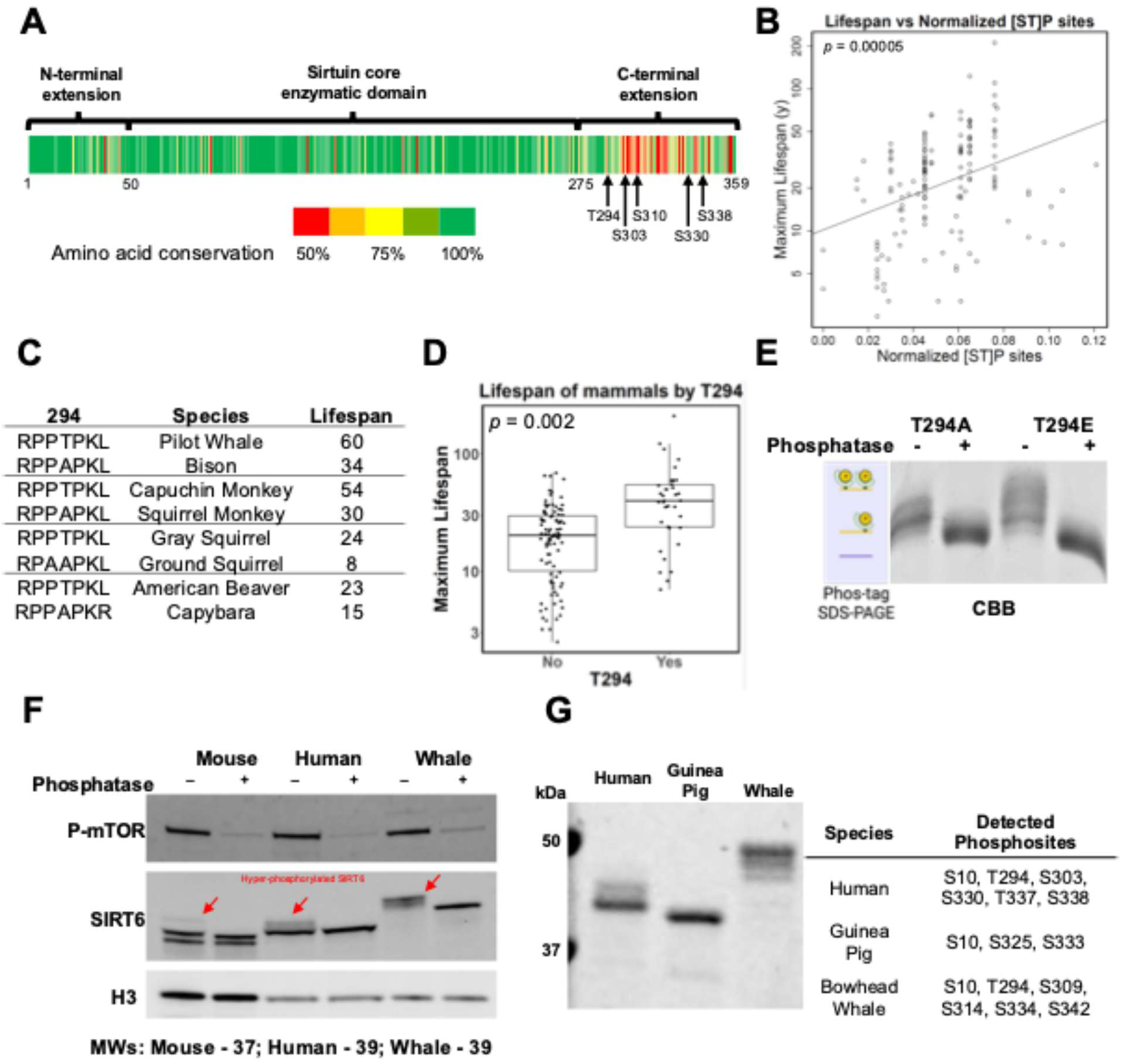
SIRT6 hyperphosphorylation correlates with maximum lifespan. **A)** Heatmap of SIRT6 amino acid conservation across more than 150 placental mammals. Green is more conserved, and red is less conserved. Absolute conservation was computed to generate the heatmap; chemically similar amino acids are not considered conserved. **B)** Plot of species maximum lifespan against the number of predicted phosphorylation sites using the [ST]P motif, normalized to C-terminal domain length, across 150 mammals. **C)** Closely related species with large differences in lifespan and the presence or absence of phospho-acceptor site T294. **D)** Box plot of species maximum lifespan for 150 mammals separated by the presence or absence of threonine at the 294 position of SIRT6. Statistics: Both variables were correlated with lifespan using the phylogenetic generalized least squares method and had p < 0.05. **E)** Coomassie stained gel after phos-gel of recombinant SIRT6 proteins from human T294A or T294E species produced in HEK293T cells. **F)** Western blots on cell lysates from primary fibroblast cell cultures from three species (M. musculus, H. sapiens, B. mysticetus) with or without phosphatase treatment. **G)** Coomassie stained gel after SDS-PAGE of recombinant SIRT6 proteins from three species produced in HEK293T cells and corresponding MS-IDs.

We also employed this methodology to study a particular site, T294. This site is unique in SIRT6 because it is the only C-terminal site present in the well-conserved portion of the SIRT6 coding sequence, which enables it to be directly compared across species in a way not possible for other C-terminal sites. Direct observation of the aligned sequences shows that the more common residue at this position is alanine, with only a few lineages possessing a phosphoacceptor site. Amazingly, the groups that have this mutation are extremely long-lived; the broadest groups are *Haplorhini* and *Cetacea*, which are two of the longest-living mammalian clades (Figure 2C). Thus, using a predicted search for phosphorylations on the C-terminus of SIRT6 across more than 150 mammals, we find that the number of sites generally, and T294 specifically, are associated with extended longevity (Figure 2D). Running purified SIRT6 T294A or T294E recombinant human proteins on phos-gels revealed that the presence results in further phospho-modifications of the C-terminus (Figure 2E). Consistently with our bioinformatic analysis, SIRT6 from longer-lived species from diverse mammalian taxa showed a greater shift after phosphatase treatment (Figures 2F). This suggests that long-lived species have more phosphorylations on SIRT6 than short-lived species, possibly due to sequence divergence in their C-terminal domains. Interestingly, we find no difference in the mRNA expression levels of SIRT6 across species’ fibroblasts (Figure S2A). Instead, through immunoprecipitation of endogenous SIRT6 from metabolically labeled fibroblasts, we find significant differences in SIRT6 protein turnover rate between human and mouse (Figures S2B and S2C).

To validate these findings, we purified recombinant SIRT6 using our mammalian system from three species with varying numbers of predicted C-terminal phosphosites: guinea pig (2), human and bowhead whale (6). These proteins showed similar phosphorylation-associated shifts on SDS-PAGE to those observed in primary fibroblast lysates (Figure 2G). Using MS, we observed all the expected phosphorylations for these sequences based on the [ST]P pattern. The phosphopeptides were abundant in all cases where the unmodified peptide was also detectable. In summary, we find that SIRT6 C-terminal phosphorylation is highly variable across species and is positively correlated with species maximum lifespan.

### Multiple CMGC kinases can progressively phosphorylate SIRT6

We next attempted to identify the kinases and cellular conditions responsible for phosphorylation of SIRT6 C-terminus. Because many of these sites are defined by the presence of proline in the -1 or +1 positions, we focused our search within the CMGC family of proline-directed kinases. The CMGC family is a large and diverse group of kinases that play critical roles in a variety of cellular processes, including control of the cell cycle, transcription, and signal transduction pathways, and its members share a near-requirement for proline in the +1 position [33–35]. We found that both a MAPK and a HIPK could phosphorylate SIRT6 *in vitro* as measured by electromobility shift on SDS-PAGE, while a CDK-like kinase (CLK) was substantially less effective (Figure 3A). We then treated human fibroblasts with inhibitors of CMGC kinase family members [36–38]. Inhibition of the MAPK, GSK, and CDK families did not produce a noticeable effect on the gel migration of SIRT6 (Figure S3A), while inhibition of the DYRK kinases, a part of CMGC family, substantially reduced the upper SIRT6 doublet band (Figure 3B). These results suggest that the DYRK kinase family, which includes HIPKs, likely contributes to the regulation of SIRT6 C-terminal phosphorylation.

**Figure 3.**
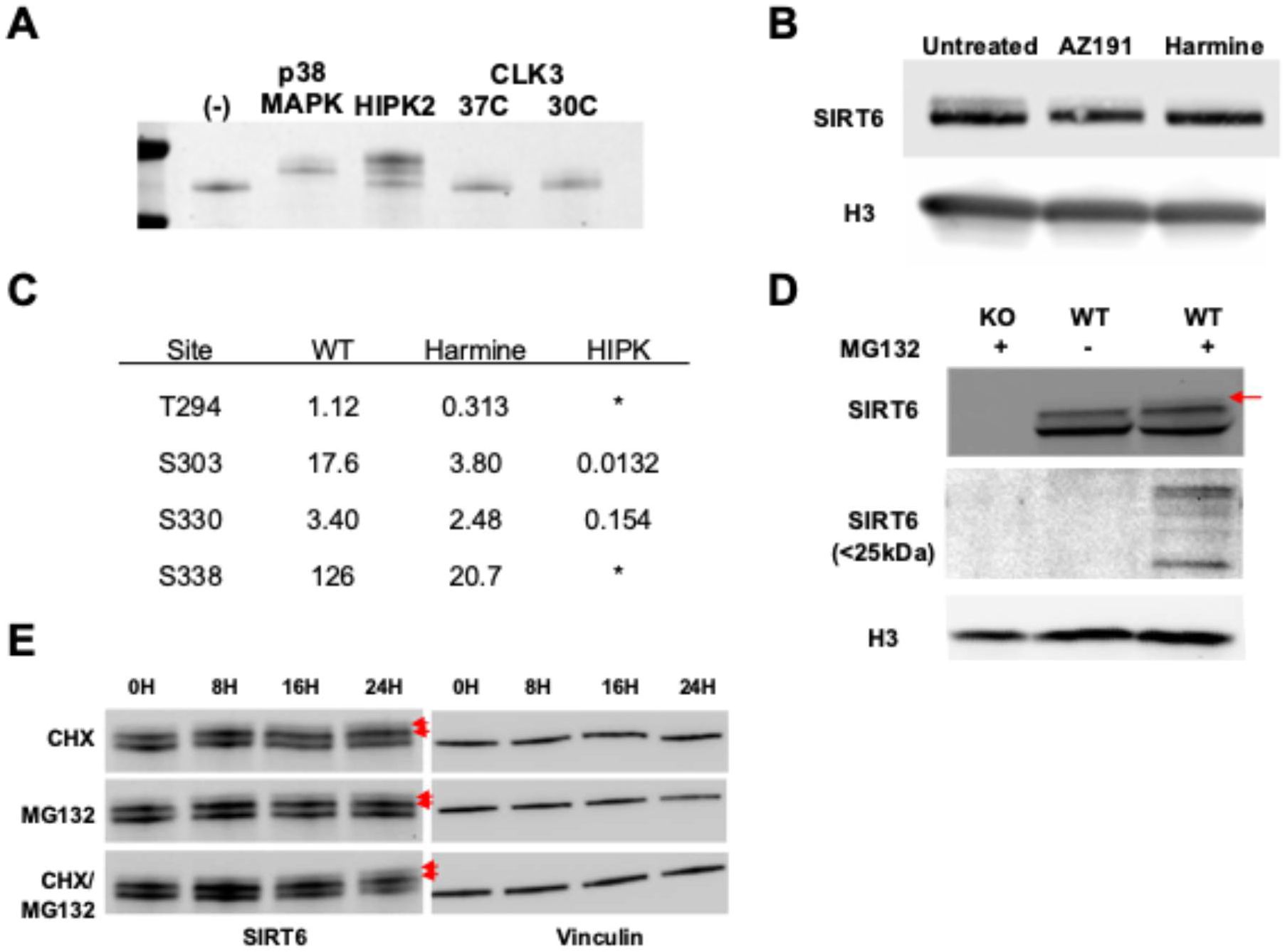
Multiple CMGC kinases progressively phosphorylate SIRT6. **A)** Coomassie stained gel after SDS-PAGE of human WT SIRT6 purified from E. coli and incubated with various proline-directed kinases in kinase buffer for one hour. **B)** Western blots of fibroblast cell lysates after 24h treatment with DYRK inhibitors. **C)** Table of relative phosphopeptide intensities from LC-MS/MS on recombinant human SIRT6 purified from HEK293 cells under normal and HIPK inhibited conditions and compared with SIRT6 purified from E. coli and treated with recombinant HIPK. The T294 and S338 unmodified peptides were below the threshold of detection, rendering the relative intensity as an estimate. For both (C), relative intensity is computed by dividing the precursor ion intensity of the phosphopeptide by that of the same peptide without the phosphorylation. These numbers are not comparable vertically in this table, only horizontally. **D)** Western blot from primary human fibroblasts or SIRT6 KO fibroblasts treated with or without 10uM MG132. **E)** Western blot from HEK293T cells treated with cycloheximide, MG132, or both for the indicated time periods.

We then sought to identify the specific phosphosites that were affected by the activity of DYRK kinases. We produced SIRT6 recombinantly in HEK293 cells in the presence or absence of the DYRK inhibitor and performed MS on the purified proteins. We also included the *in vitro* HIPK-treated sample from Figure 3A as comparison. The results showed that both T294 and S338 are under control of the DYRK family, as these sites were responsive both to inhibition and *in vitro* phosphorylation (Figure 3C). DYRK family members have been implicated in many cellular processes, including RNA splicing, DNA repair, and proteostasis, and appear to be activated by a variety of conditions [39–43]. Thus, we treated fibroblasts with a range of stresses, including oxidative stress, topoisomerase poisoning, nutrient stress, and temperature shock (Figure S3B). We found that none of these conditions were sufficient to produce a visible change to electromobility like that observed after DYRK inhibition, suggesting that the control of SIRT6 C-terminal phosphorylation is insensitive to these stressors. These results suggest that SIRT6 C-terminal phosphorylation is performed by DYRK family kinases constitutively in cell culture.

We were able to identify an increase in endogenous phosphorylated SIRT6 in two unique and seemingly synergistic cases: proteasome inhibition by MG132 and translation inhibition by cycloheximide. A 24-hour treatment with MG132 led only to a slight accumulation of total SIRT6 protein in human fibroblasts but surprisingly lead to the appearance of a third further phosphorylated proteoform (Figure 3D). We also identified the appearance of lower molecular weight SIRT6-specific bands between 20 and 27 kDa upon proteasomal inhibition (Figures 3D and S3C). We speculate that these are rapidly degraded fragments of SIRT6, though it is presently unclear how these fragments are formed. Surprisingly, treatment with MG132, cycloheximide, or both in confluent HEK293 cells and quiescent human fibroblasts overexpressing SIRT6 resulted in clear accumulation of all phosphorylated proteoforms over time (Figures 3E and S3D). This result suggests that SIRT6 may be progressively phosphorylated over the course of its molecular lifetime.

### Phosphorylation of SIRT6 enhances oxidative stress resistance via PARP1

Next, we turned our attention to the effects of C-terminal phosphorylation on SIRT6 interactions. Mass spectrometry analysis of SIRT6 purified from HEK293 showed that several known SIRT6 interacting proteins, including core histones, Ku70/80, and PARP1, consistently co-purify with SIRT6 (not shown). We then confirmed this observation by running human and mouse SIRT6 purified from HEK293 on Western blot (Figure 4A). Interestingly, much lower levels of PARP1 co-purified with mouse SIRT6 compared to human SIRT6.

**Figure 4.**
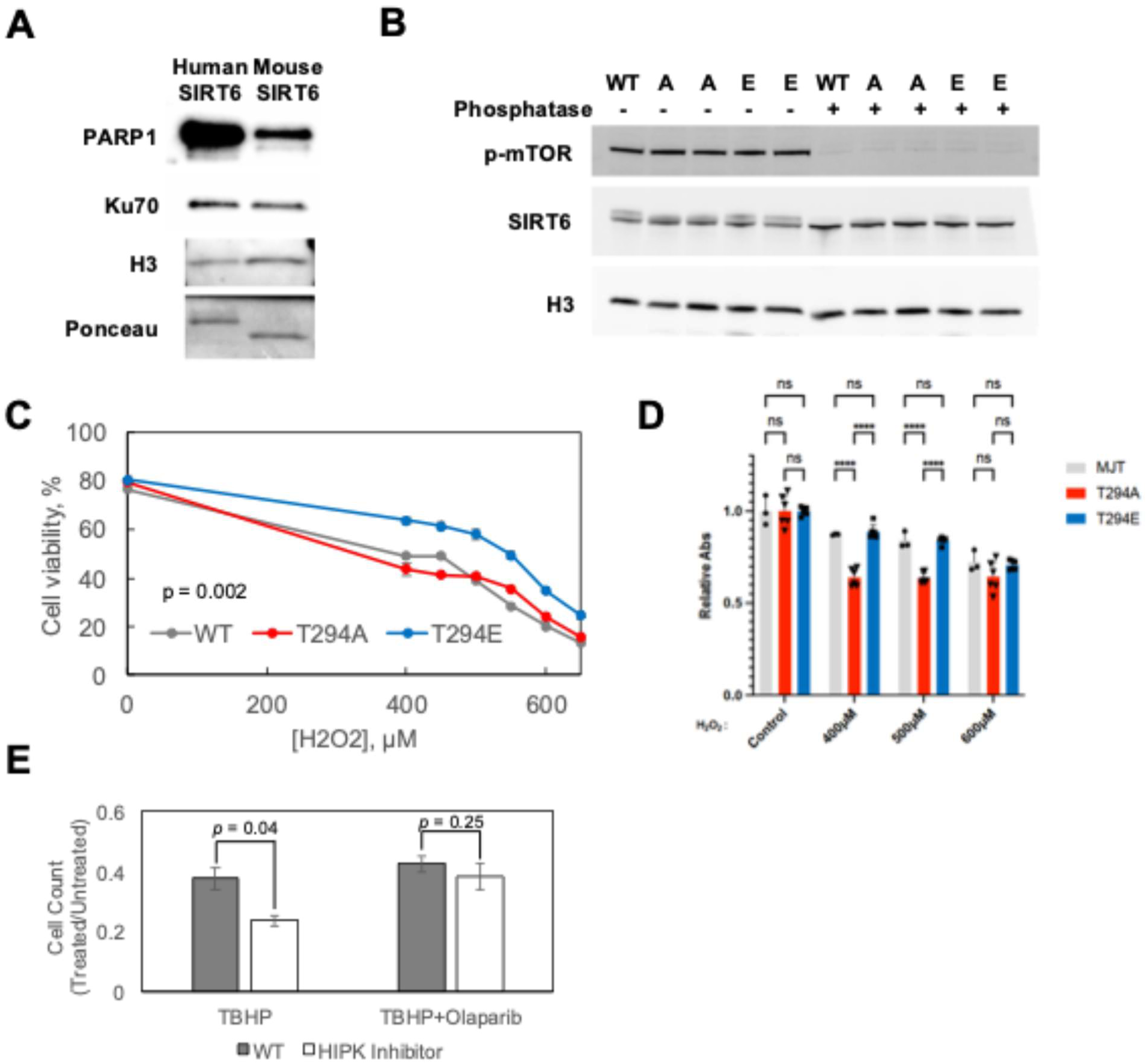
Hyperphosphorylation enhances oxidative stress resistance via PARP1. **A)** Western blots for co-purifying proteins on recombinant human or mouse SIRT6 preps produced in human cells. Ponceau image is the SIRT6 band at ∼40kDa. **B)** Western blots of cell lysates with or without phosphatase from human fibroblast cell lines harboring homozygous mutant endogenous SIRT6 alleles for WT, T294A “A”, or T294E “E” SIRT6. Replicate genotypes are independent clones. **C)** Cell viability of T294 mutant fibroblasts as measured by Annexin V and propidium iodide negativity five days after treatment with various concentrations of hydrogen peroxide. p < 0.05 for the comparison of T294E to the WT and T294A genotypes as measured by two-way ANOVA and Tukey’s HSD. **D)** WST-1 viability assay comparing the WT, T294A, & T294E genotypes. **** = p < 0.0001 by Tukey’s HSD as follow-up to two-way ANOVA. **E)** Cell number relative to untreated cells 96 hours after treatment with 100µM tert-butyl hydrogen peroxide (TBHP) or TBHP + 100nM Olaparib. Harmine = WT fibroblasts pre-treated with harmine for 8 hours prior to TBHP.

Human SIRT6 contains a phosphorylation site at position 294 (T294) which is lacking in the mouse SIRT6. Remarkably, human SIRT6 T294A mutant showed reduced PARP1 co-purification as measured by Western blot (Figure S4A). There was no effect on Ku70/80 co-purification by either T295A or T294E mutation. Additionally, general kinase inhibition during protein production with the promiscuous kinase inhibitor dorsomorphin produced the same reduction in PARP1 co-purification as the T294A mutation (Figure S4A). Interestingly, kinase inhibition had the same effect on PARP1 co-purification for WT SIRT6 and the T294E mutant, while it had no effect on the T294A mutant. These results suggest that T294 phosphorylation facilitates PARP1 interaction but is not essential for it.

We then tested if the changes in PARP1 interaction resulted in alterations to cellular function. Because T294 mutation alone was sufficient to affect the interaction with PARP1, we used CRISPR-Cas9 to generate clones with homozygous knock-in T294A and T294E alleles at the endogenous locus of h-TERT immortalized diploid fibroblasts (Figure S4B). We observed the expected changes in SIRT6 SDS-PAGE electromobility and no significant alterations in SIRT6 protein expression (Figures 4B and S4C).

Then, we tested the resistance of the mutated cell lines to oxidative damage, as PARP1 is particularly important for the repair of ssDNA breaks induced by oxidative lesions. Measuring survival after hydrogen peroxide treatment demonstrated increased survival of the SIRT6 T294E cells, consistent with the hypothesis of increased PARP1 activity and subsequent repair of oxidative DNA lesions in these cells (Figure 4C). WST-1 cell viability assay confirmed a trend of reduced oxidative stress resistance in the T294A mutant (Figure 4D). Inhibition of HIPK using harmine treatment also caused a decrease in survival to tert-butyl hydrogen peroxide in a manner dependent on PARP1 (Figure 4E). Together, the results for the mutant genotypes are consistent with the hypothesis that T294 phosphorylation enhances oxidative stress resistance in a PARP1-dependent manner.

### Molecular modeling predicts that C-terminal SIRT6 phosphorylation enhances interaction with PARP1

To predict the potential effects of SIRT6 hyperphosphorylation on its structure and interactions, we used the publicly available AlphaFold3 multimer server to predict the structures of human SIRT6 with and without known N- and C-terminal phosphorylations as monomers, homodimers, and heterodimers in complex with human PARP1 [44]. In all instances, we find that with phosphorylation of S10, the C-terminus, or both, AlphaFold predicts a more structured C-terminus, and overall protein, than the endogenous intrinsically disordered termini (Figures 5A-C). All SIRT6 homodimers also show increased structure in these regions with significant enhancement of the intricacy of the predicted interaction between SIRT6 molecules (Figures S5A-D). This trend persisted for the SIRT6-PARP1 heterodimers. When unmodified, the SIRT6 C-terminus is unstructured and is not in proximity to either SIRT6 itself or PARP1 (Figure 5A). However, upon phosphorylation, the SIRT6 C-terminus is predicted to fit tightly in the known DNA binding pocket of PARP1 (Figure 5B). We find that this binding pocket is positively charged and free from any overt steric clashes or hindrances (Figure 5C). Thus, we propose that SIRT6 C-terminal hyperphosphorylation may function to tether SIRT6 to PARP1 while still allowing enough residual flexibility for SIRT6 to activate PARP1 via ribosylation at K521.

**Figure 5.**
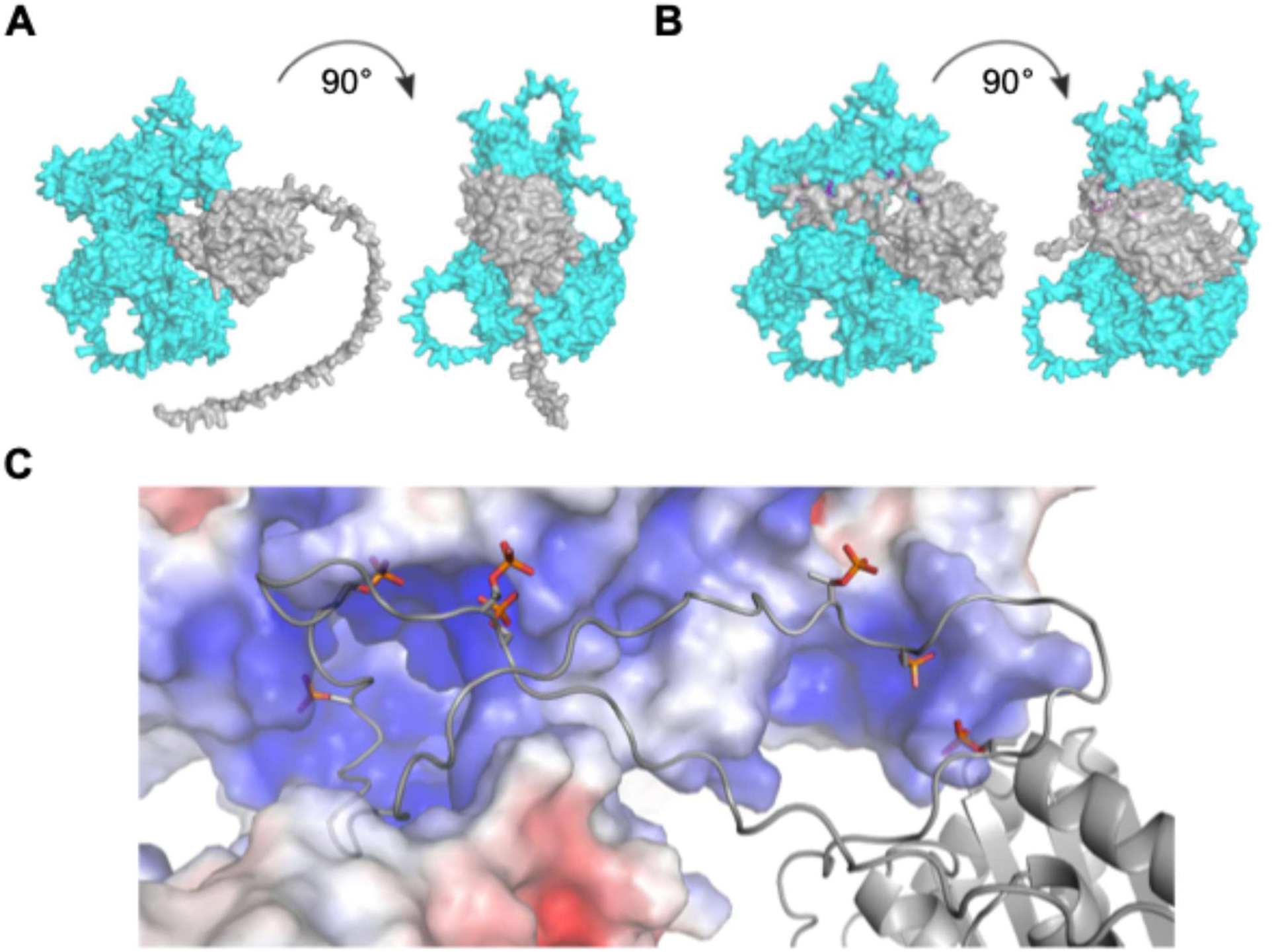
Hyperphosphorylation is predicted to enhance PARP1 interaction. **A)** AlphaFold3 structures of unmodified SIRT6 as heterodimer with PARP1. Cyan = PARP1, Gray = SIRT6. **B)** AlphaFold3 structures of P7 SIRT6 as heterodimer with PARP1. P7 = seven C-terminal phosphorylations (T294, S303, T305, S326, S330, T337, S338) **C)** Detailed view of steric clash, local charge, and putative interaction interface between the phosphorylated C-terminus of SIRT6 (foreground, wireframe model) and a basic region of PARP1 (background, space-filing model).

In summary, we show that the number of SIRT6 C-terminal phosphorylation sites correlates positively with species’ maximum lifespan. Furthermore, our results suggest that C-terminal phosphorylation promotes SIRT6 interaction with PARP1 and resistance to oxidative stress.

## DISCUSSION

SIRT6 is a well-studied longevity factor that has generated excitement for its potential in anti-aging therapeutic strategies. Despite extensive literature documenting its roles in aging and disease, relatively little is known about its post-translational regulation. Here, we present evidence that phosphorylation of the SIRT6 C-terminal tail is abundant, positively correlated with lifespan, and improves the interaction between SIRT6 and PARP1. We show that major phosphorylation sites on SIRT6 share a proline-directed motif, and that this motif is conserved even across species with significantly divergent C-terminal amino acid sequences. The number of these sites on the C-terminus of SIRT6 is also positively associated with lifespan. In human cells, we identified at least 4 distinct phospho-modified SIRT6 proteoforms. We find that SIRT6 C-terminal phosphorylation can be carried out by multiple kinase families.

Our data suggest that several proline-directed kinases can efficiently phosphorylate multiple sites on SIRT6. The lack of kinase specificity of these sites is consistent may underscore the complex regulation of this phosphorylation. It is likely that there is substantial co-regulation of these sites. Common regulation by multiple PTMs is indeed a frequent feature of intrinsically disordered regions, such as the SIRT6 C-terminus [45, 46]. It would be interesting to measure SIRT6 phosphorylation in different cell and tissue types, which may identify divergent phosphorylation patterns.

We provide evidence that SIRT6 phosphorylation is in part influenced by DYRK family activity. This family of kinases has diverse roles in transcriptional regulation, differentiation, DNA repair, and stress response [37, 39–43, 47], a feature they share with SIRT6. Future studies will elucidate the extent to which the relationship between DYRKs and SIRT6 influences development and stem cell aging.

We found that C-terminal phosphorylation enhances interaction with PARP1 and improves resistance to oxidative damage in a PARP1-dependent manner. SIRT6 is known to be a rapid responder to SSBs along with PARP1, and its absence delays the recruitment of several required BER proteins [23, 48]. Thus, these results offer the exciting possibility that SIRT6 C-terminal phosphorylation enhances this role, and that this enhancement conferred sufficient fitness benefit to be positively selected during the evolution of longevity.

## MATERIALS & METHODS

### Cell Culture

Cell culture was performed with standard techniques. Primary fibroblasts were cultured in Eagle’s Minimum Essential Medium supplemented with 15% FBS and pen/strep and transformed cell lines were cultured in Dulbecco’s Modified Eagle’s Medium with 10% serum and pen/strep. All cultures were grown at 37°C in a humidified atmosphere with 5% CO_2_ and 4% O_2_.

### Generation of cell lines

Cell culture was primarily performed on derivatives of an immortalized HCA2 human cell line “MJT”. T294 knock-in mutants were generated using CRISPR-Cas9. A guide RNA targeting a cut just 3’ of T294 was designed using IDT’s design tool; crRNA sequence CTCCTTGGGCTCCAGCTTGG. 200bp HDR donor sequences were designed to incorporate the desired mutation into the repaired chromosome, along with two silent mutations in the crRNA target sequence to discourage further cutting after successful HDR. All reagents were obtained from IDT. Duplexes were formed by mixing crRNA and tracrRNA 1:1 and heating at 95°C for 5min and cooling on benchtop for 5min. Ribonucleoprotein (RNP) complexes were formed by mixing 104 pmol of HiFi Cas9 and 120 pmol RNA duplex and incubating on benchtop for 20 min. RNP delivery was performed via nucleofection with a Lonza 2b nucleofection system in the presence of 4µM IDT electroporation enhancer and 4 µM HDR donor oligo. Transfected cells were cultured overnight with 1µM HDR enhancer after which the enhancer was removed. Two days after transfection cells were replated in 96-well formats at 0.5 cells/well to form clonal populations. Clones were screened both via PCR with primers overlapping the mutation site as well as Sanger sequencing. PCR primer sequences were as follows: Common forward primer CTCCTTGGGCTCCAGCTTGG, WT reverse primer GCTCCAGCTTCGGGGT, T294A reverse primer GCTCCAGCTTCGGGGC, T294E reverse primer GCTCCAGCTTCGGCTC.

### Drug treatments

All drug treatments were performed in 6-well plates 24 hours after plating 200,000 cells unless otherwise specified. Etoposide was used at 10µM for 48 hours, while all other drug treatments were for 24 hours. Concentrations were as follows: MG132 10 μM (Millipore Sigma), harmine hydrochloride 10 μM (Selleckchem cat#S3817), dorsomorphin 5μM (Selleck cat#S7840), SB202190 5 µM (Selleck cat#S1077), lithium chloride 50 mM, dinaciclib 100 nM (Selleck cat#S2768), paraquat 1 mM (Sigma cat#856177), 2-deoxyglucose 5 mM (Sigma cat#D4601).

### Oxidative stress resistance assay

Cells were plated in 12-well plates at a density of 30,000 cells/well and allowed to attach overnight. The harmine condition were WT cells pre-treated with 10 μM harmine hydrochloride for 8 hours beginning the morning after plating. Then, cells were exposed to 250µM tert-butyl hydrogen peroxide (TBHP) for one hour with or without 100 nM Olaparib. Olaparib or harmine treatments were continued for 6 hours post-removal of TBHP to ensure repair took place in their presence before replacement of normal medium. Four days later cells were collected and counted.

### WST-1 Cell Viability Assay

Cells expressing different SIRT6 variants were seeded in 96-well plates at 5,000 cells per well and allowed to adhere overnight. The following day, media containing 0, 400, 500, or 600 µM H₂O₂ were applied. After 48 hours of incubation, 10 µl (per well) of Cell Proliferation Reagent WST-1 (Cellpro-ro, Merck) was added and incubated for 2 hours. Plates were briefly shaken (1 minute), and absorbance was measured using a plate reader at 440 nm (formazan product) with 600 nm as reference. Absorbance values were background-corrected (using cell-free wells) and normalized to the average of untreated control. Each condition was tested in triplicate. Both SIRT6 T294E and SIRT6 T294A were tested in two biological replicate cell lines.

### Immunoprecipitation

Immunoprecipitation (IP) was performed using the Dynabeads Protein A IP kit (Invitrogen, Cat#1006D) according to the manufacturer protocol. Briefly, magnetic beads were incubated with the appropriate IgG in Antibody Binding Buffer for 10min, washed once, and resuspended in 1 mL of cell lysate. Binding was performed overnight at 4°C. The beads were washed 3x with kit Wash Buffer and eluted in 20 μL of kit Elution Buffer, which was then neutralized with 5 μL of 1M Tris pH 7.5.

### Western blot

Blots were performed according to standard methods using a semi-dry transfer system and PVDF membranes blocked with 5% nonfat dry milk. Antibodies used are as follows: SIRT6 CST cat#12486, H3 CST cat#4499, p-mTOR CST cat#5536, PARP1 Thermo cat#13371-1-AP, Ku70 Abcam cat#ab202022, p-Rb CST cat#9308S, γ-H2AX Millipore cat#05-636, Cyclin B1 CST cat#4138, Vinculin CST cat#4650.

### Phosphatase treatment

40 μL of concentration-normalized protein samples were incubated with 5µL of 10X phosphatase buffer, 5 μL of 10 mM MnCl_2_, and 1 µL of lambda protein phosphatase (NEB cat#P0753L) at 30°C for 30min.

### Prokaryotic purification system

Coding sequences were cloned into pET11a vectors and transformed into *BL21 DE3* strain *E. coli*. Cultures were grown in LB broth supplemented with 50 μg/mL ampicillin at 37°C in a shaking incubator until the culture optical density reached 1.0, when 0.4 mM IPTG was spiked in and the temperature was reduced to 25°C. After 18h induction cells were pelleted at 6,000xg. The pellet was resuspended in purification buffer (300 mM NaCl, 50 mM Tris pH=7.5, 10mM imidazole, 5% glycerol, and Pierce protease inhibitors) at 4 mL/g wet mass. 20mg of egg white lysozyme was added and the tube was rotated for 1h at 4°C before sonication 6x 20s at 50% power, resting on ice in between cycles. After lysis the sample was centrifuged at 20,000xg for 1h at 4°C and the supernatant removed. 1mL packed volume equivalent of Ni-NTA agarose resin (Qiagen Cat#30250) was added to the tube and rotated at 4°C for 2h. The resin was centrifuged at 1,500xg for 5min and resuspended in 10 volumes of wash buffer (300 mM NaCl, 50 mM Tris pH=7.5, 5% glycerol, 30mM imidazole) a total of 5 times. The sample was the eluted in wash buffer + 500 mM imidazole and dialyzed overnight in storage buffer (150 mM NaCl, 20 mM Tris pH 7.5, 20% glycerol, 1 mM DTT) with one buffer change. Protein concentration was assessed with a BCA assay and purity was verified with SDS-PAGE.

### Mammalian purification system

Coding sequences were cloned into pFN21A vectors (Promega Cat#G2821) and transfected into HEK293T cells using Viafect (Promega Cat#E4982) at 25 µg DNA/15cm plate, with two plates per condition. Cells were harvested 72 hours after transfection using a cell scraper after an ice-cold PBS wash and duplicate plates were pooled. Cells were pelleted at 300xg for 5min and flash frozen in liquid nitrogen. Protein purification was performed using the HaloTag mammalian purification kit (Promega Cat#G6790) according to the kit protocol. In brief, pellets were resuspended in 1 mL of lysis buffer (150 mM NaCl, 50 mM Tris pH 7.5, 1% Triton X-100, and protease inhibitors) and digested with 20 µL DNase I for 15 min at 25°C. Lysates were diluted with purification buffer (150 mM NaCl, 20 mM Tris pH 7.5, 1mM DTT, 0.005% CA-135) and centrifuged at 15,000xg for 30min at 4°C, after which the supernatant was added to 50 µL packed resin volume of HaloTag resin and incubated overnight at 4°C. Resin was washed 4x in purification buffer and eluted twice in 150 µL using HaloTEV protease. Protein concentration was measured using a BCA assay and purity verified with SDS-PAGE.

### In vitro kinase assay

One μg of prokaryotic-derived recombinant human SIRT6 was prepared with 0.1μg recombinant active kinase from Abcam (HIPK2: ab268639; MAPK: ab268832; CLK3: ab85759) in kinase buffer (10 mM MgCl_2_, 20 mM tris pH 7.5, 1 mM DTT). Samples were incubated at 37°C for 10 min before being spiked with 1mM ATP and incubated for 1h.

### Targeted dSILAC of SIRT6

Targeted dSILAC on endogenous SIRT6 from primary mouse or human fibroblasts was performed as follows. Individual 15 cm plates of quiescent cells were adapted to dialyzed FBS over a period of 4 days. Labeling was then performed on each plate with ^13^C_6_ arginine & ^13^C_6_ lysine for 0 hours, 6 hours, 12 hours, 24 hours, 48 hours, 72 hours, or 144 hours. Immunoprecipitation was then conducted as described above. In-gel digests were then conducted as described below.

### Mass spectrometry

Samples were prepared using S-Traps (Protifi Cat#CO2-micro-80) essentially according to manufacturer protocol. In brief, samples were denatured with one volume of denaturing buffer (10% SDS, 100 mM TEAB), reduced with 10 mM DTT at 55°C for 30 min, and alkylated with 15 mM iodoacetamide at 25°C in the dark for 30 min. Samples were then acidified with 27.5% phosphoric acid and diluted in wash buffer (90% methanol, 100 mM TEAB). They were loaded onto the S-trap column and washed 5x with wash buffer. Trypsin digestion was performed on-column overnight with 2 µg of trypsin in 40 µL of 50 mM TEAB at 37°C before sequential elution with 50 mM TEAB, 0.2% formic acid (FA)/5% acetonitrile (ACN), and 50% ACN and evaporation with a speedvac. Peptides were resuspended in 0.2% FA and 5% ACN at 0.5 mg/mL.

### In-gel digestion

Gel was washed in distilled water overnight at room temperature. Individual bands were cut and diced into ∼1mm^3^ pieces and placed into lo-bind tubes. Gel pieces were washed with 100 µl of 50 mM ammonium bicarbonate (ABC) for 10 minutes, follow by washing twice with 100 µl of 25 mM ABC in 50% acetonitrile (ACN) for 10 minutes, shaking continuously. Gel pieces were further dehydrated by washing with 100% ACN for 5 minutes while shaking. Excess ACN was removed by drying in a speed vac (Labconco). Disulfide bonds were reduced by adding 100 µl of 10mM dithiothreitol (DTT, Sigma) in 50mM ABC and incubated at 55°C for 1hr. Alkylation was performed by adding 100 µl iodoacetamide (IAA, Sigma) in 50 mM ABC and incubated at RT for 30 minutes in darkness. Gel pieces were washed of excess DTT and IAA with 100 µl of 50 mM ABC for 10 minutes followed by 100 µl of 25 mM ABC in 50% ACN for 10 minutes, continuously shaking. Gel pieces were further dehydrated by washing with 100 µl of 100% ACN for 5 minutes while shaking. Excess ACN was removed by drying in a speed vac. 500 ng of trypsin (Thermo), AspN (Thermo), or chymotrypsin (Promega) in 50 mM ABC were added to the dehydrated gel pieces to completely cover and incubated for 1 hr at RT to allow gels to fully rehydrate followed by overnight incubation at 37°C. Supernatant containing digested peptides was transferred to clean lo-bind tubes. Peptides were further extracted from the gel pieces by washing twice with 50/50 ACN/0.1% TFA for 25 minutes followed by washing wit 50 µl 100% ACN for 1 minute, continuously shaking. Pooled extracts were frozen and dried down in a speed vac. Samples were subsequently desalted using homemade C18 spin columns and sample were frozen and dried down in a speed vac.

### LC-MS/MS analysis

#### Astral data collection

Peptides were injected onto an Aurora Elite TS 15cm C18 column (IonOpticks) heated to 40°C, with a Vanquish Neo HPLC (Thermo Fisher), connected to a Orbitrap Astral mass spectrometer (Thermo Fisher). Solvent A was 0.1% formic acid in water while solvent B was 0.1% formic acid in 80% ACN. Ions were introduced using a, EasySpray source operating at 2 kV. The gradient began at 1% B, increased to 5% B in 0.1 minutes, then increased to 30% B over 11.3 minutes, then increased to 40% B over 1.5 minutes, then increased to 99% B over 0.1 minutes and was held for 2 minutes for a total run time of 14 minutes before equilibrating the column to starting conditions for 1.5 minutes. The Orbitrap Astral was operated in data-dependent mode with MS1 scans acquired in the Orbitrap and MS2 scans acquired in the Astral. The cycle time was set to 0.5s. Monoisotopic precursor selection was set to “peptide”. The full scan was performed over a range of 375-1,400 m/z with a resolution of 120,000 at m/z 200, an AGC target of 300%, and a maximum injection time of 10ms. Peptides with a charge state between 2 and 5 were selected for fragmentation with a minimum intensity filter set to 5e3. Precursor ions were fragmented by higher energy dissociation (HCD) using a collision energy of 27% with an isolation width of 2 m/z. The MS2 scan was performed over a range of 140-2,000 m/z with an AGC target of 200% and a maximum injection time of 10 ms. Dynamic exclusion was set to 5s and to exclude after 1 time using a mass tolerance of +/- 5 ppm with exclude isotopes set to “True”.

#### Lumos data collection

Peptides were injected onto a homemade 30 cm C18 column with 1.8 µm beads (Sepax), with an Easy nLC-1200 HPLC (Thermo Fisher), connected to a Fusion Lumos Tribrid mass spectrometer (Thermo Fisher). Solvent A was 0.1% formic acid in water, while solvent B was 0.1% formic acid in 80% acetonitrile. Ions were introduced to the mass spectrometer using a Nanospray Flex source operating at 2 kV. The gradient began at 3% B where it was held for 2 minutes, increased to 10% B in 5 minutes, then increased to 38% B over 38 minutes, then increased to 90% B over 3 minutes where it was held for an additional 3 minutes, before returning to starting conditions over 2 minutes and re-equilibrating for 7 minutes. The Fusion Lumos was operated in data-dependent mode with MS1 scans acquired in the Orbitrap and MS2 scans acquired in the ion trap. The cycle time was set to 1.5 s. Monoisotopic precursor selection was set to “peptide”. The full scan was performed over a range of 375-1,400 m/z with a resolution of 120,000 at m/z 200, an AGC target of 100%, and a maximum injection time of 50 ms. Peptides with a charge state between 2 and 5 were selected for fragmentation with a minimum intensity filter set to 1e4. Precursor ions were fragmented by collision-induced dissociation (CID) using a collision energy of 30% with an isolation width of 1.1 m/z. The MS2 scan was performed with an AGC target of 100% and a maximum injection time of 10ms. Dynamic exclusion was set to 20s and to exclude after 1 time using a mass tolerance of +/- 10 ppm with exclude isotopes set to “True”.

### Mass spectrometry data search

Mass spectra were searched using FragPipe version 21.1. The LFQ-phos pipeline was loaded and the default human reference proteome, with the addition of any recombinant protein sequence, was used as reference. The LFQ-phos pipeline used MSFragger [49] 4.0 for the search with methionine oxidation, cysteine alkylation, N-terminal acetylation, and STY phosphorylation set as variable mods. The machine learning tools MSBooster [50] and Percolator [51] were used to assist with peptide identification. Then, ProteinProphet [52] and PTMProphet [53] were applied to identify proteins and PTM site localization, respectively. The program used Philosopher [54] to apply a false discovery rate threshold of 0.01 to the data, and finally used IonQuant[55] version 1.10.12 to perform label-free quantitation. After the search, relative protein quantification was performed based on the unique intensity measurement in the combined_protein.tsv output file. Only proteins with a protein probability score above 0.95 as determined by ProteinProphet were considered. Relative modified peptide abundance was performed by dividing the intensity of the modified peptide by the intensity of the same unmodified peptide in the combined_modified_peptide.tsv output file. Only those peptides in which the modification localization was at least 80% confident according to PTMProphet were selected.

### Phylogenetic comparison of SIRT6 features

SIRT6 sequences were obtained from NCBI [56] and aligned using Clustal Omega [57]. Species maximum lifespans were obtained from the AnAge database [32]. The full mammalian phylogeny was downloaded from Vertlife [58]. Sequence features including predicted phosphorylation site number were counted using a custom Python script. Correlations between lifespan and sequence features were performed using the R phytools and ape packages to perform phylogenetic generalized least squares analysis (PGLS) and tree generation, respectively. PGLS was performed using the default Brownian evolutionary model on log10-transformed maximum lifespans from more than 150 mammals over the following individual sequence features: total predicted phosphorylation sites, presence of T294, net charge, proportion of charged residues, and proportion of prolines.

### AlphaFold3 multimer predictions and molecular modeling

All predictions were conducted in the AlphaFold3 online server GUI using human Uniprot protein sequences and further resolved for analysis in Pymol [59].

## AUTHOR CONTRIBUTIONS

JG, MM, EH, GT, SG, AS, & VG conceived of the experiments. JG, MM, JG, CL, EH, JCM, GT, JA, KAW, KS, JRH, PB, & VRA performed the experiments. JG, MM, KAW, KS, JRH, SG, AS, & VG analyzed data and conducted bioinformatic analyses. JG, MM, JG, AS, & VG wrote the manuscript.

## ACKNOWLEDGEMENTS

The authors would like to thank the members of the Ghaemmaghami, Gorbunova, Seluanov, and Fu laboratories for helpful discussions.

## DATA AVAILABILITY

All raw and processed MS data were uploaded to the ProteomeXchange Consortium via the PRIDE partner repository with identifier PXD076813.

## ABBREVIATIONS

ACN: acetonitrile
AGC: automatic gain control
APD: advanced peak determination
CID: collision induced dissociation
DMSO: dimethyl sulfoxide
DTT: dithiothreitol
EDTA: ethylenediaminetetraacetic acid
FDR: false discovery rate
HCD: higher energy collision dissociation
HPLC: high pressure liquid chromatography
hTERT: human telomerase
IAA: iodoacetamide
IP: immunoprecipitation
IT: ion trap
KAP1: KRAB-associated protein 1
LC-MS/MS: liquid chromatography-tandem mass spectrometry
MIPS: monoisotopic precursor selection
MS: mass spectrometry
MW: molecular weight
nLC: nanoLiquid chromatography
OT: orbitrap
PARP1: poly(ADP-ribose) polymerase 1
PD: proteome discoverer
PSM: peptide spectral match
PTM: post-translational modification
PVDF: polyvinylidene fluoride
RIPA: radioimmunoprecipitation Assay
ROS: reactive oxygen species
RT: retention time
SD: semi-dry
SIRT6: sirtuin 6
TEAB: triethylammonium bicarbonate
TFA: trifluoroacetic acid

---

**Figure S1.**
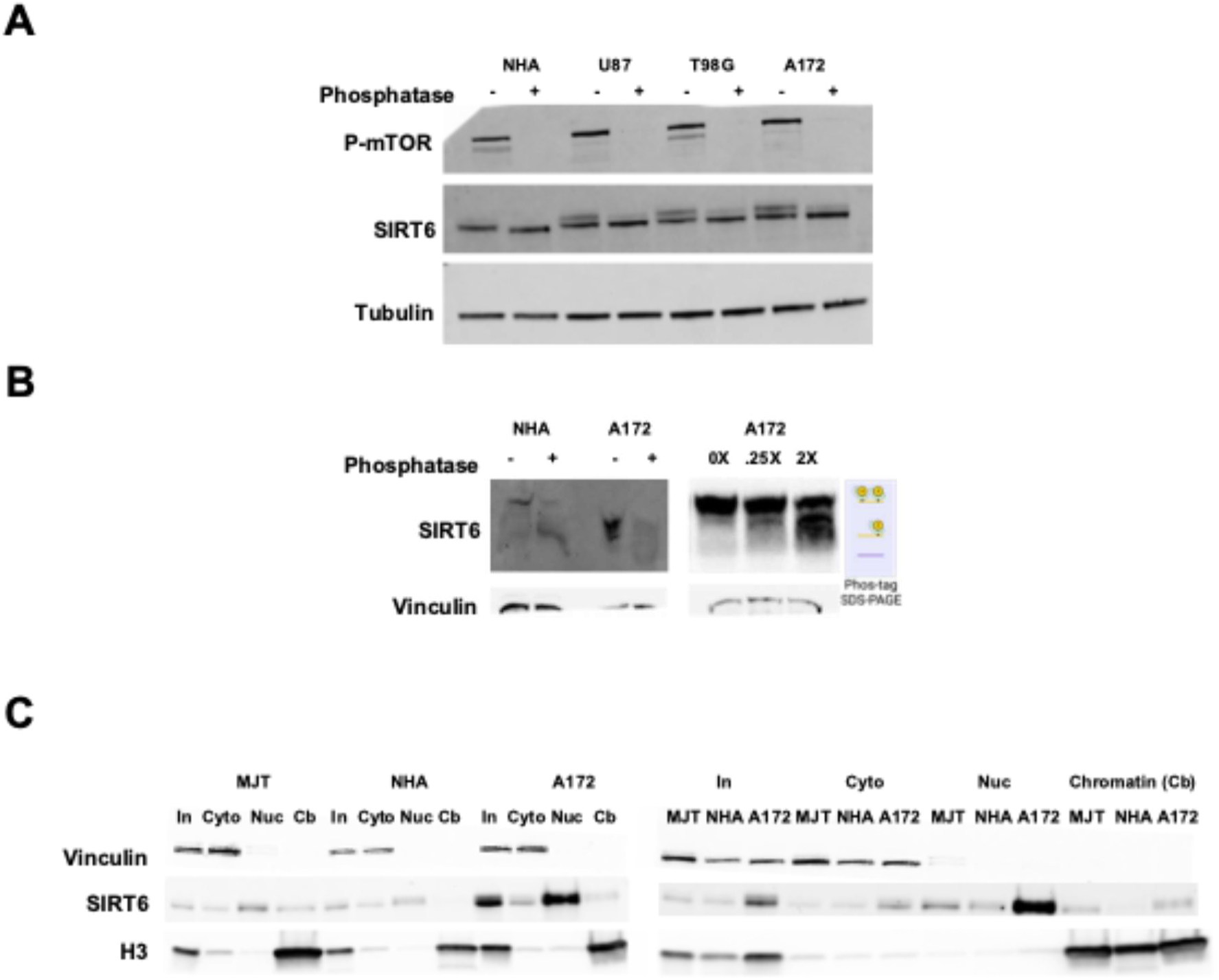
SIRT6 is hyperphosphorylated in human cells. **A)** Western blots from normal human astrocytes and three glioblastoma cell lines with or without phosphatase treatment. **B)** Western blots from NHA & A172 treated with or without phosphatase treatment and run on phos-gel. **C)** Western blots of subcellular fractionation from human fibroblasts, astrocytes, & A172.

**Figure S2.**
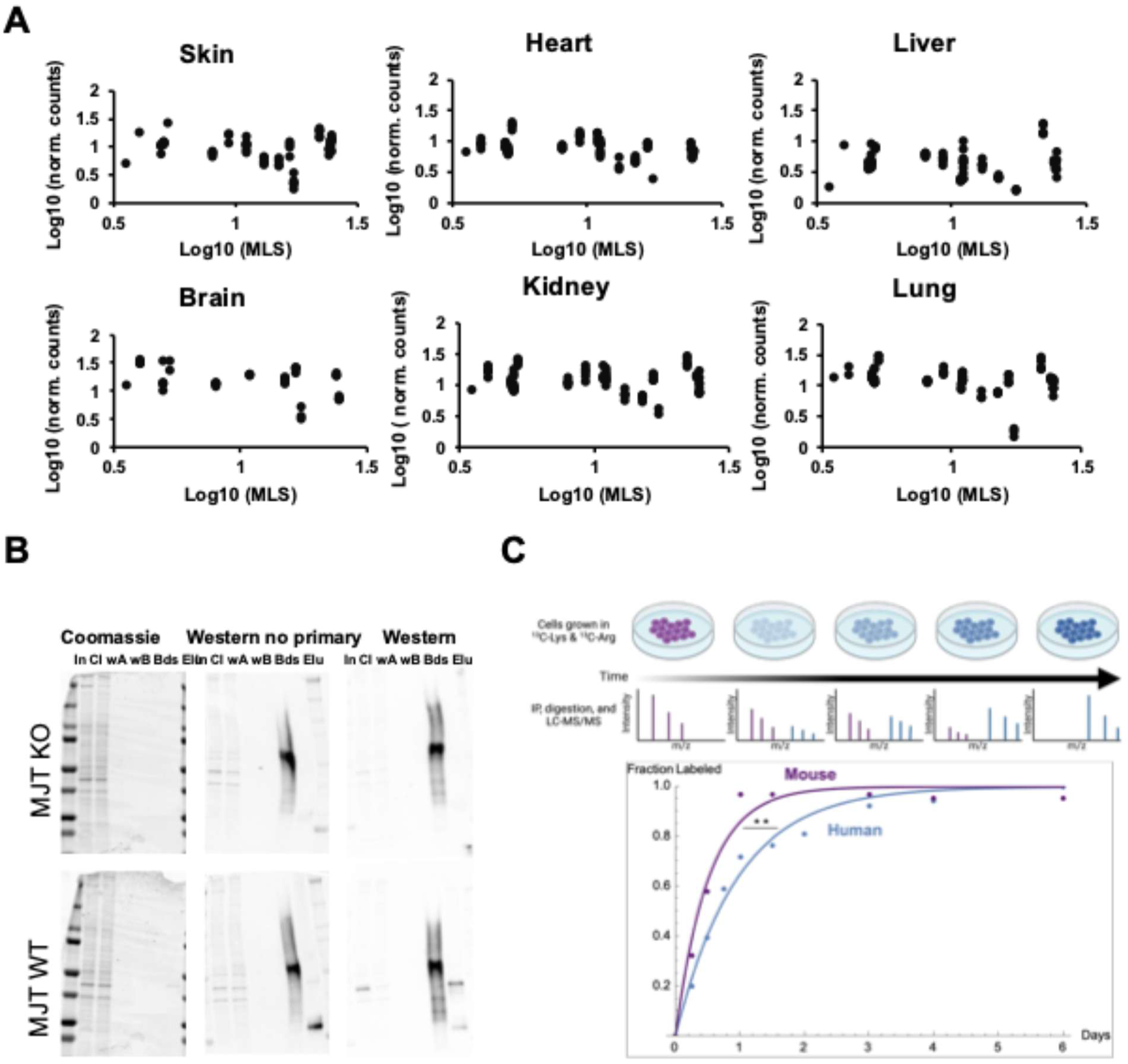
SIRT6 hyperphosphorylation correlates with maximum lifespan. **A)** Analysis of SIRT6 mRNA across species from Lu et al. 2022. **B)** Immunoprecipitation of endogenous SIRT6 from SIRTKO or WT human fibroblasts. In = input, Cl = cleared lysate, wA = wash A run-off, wB = wash B run-off, Bds = beads, Elu = elution. **C)** dSILAC of primary mouse or human fibroblasts immunoprecipitated for endogenous SIRT6.

**Figure S3.**
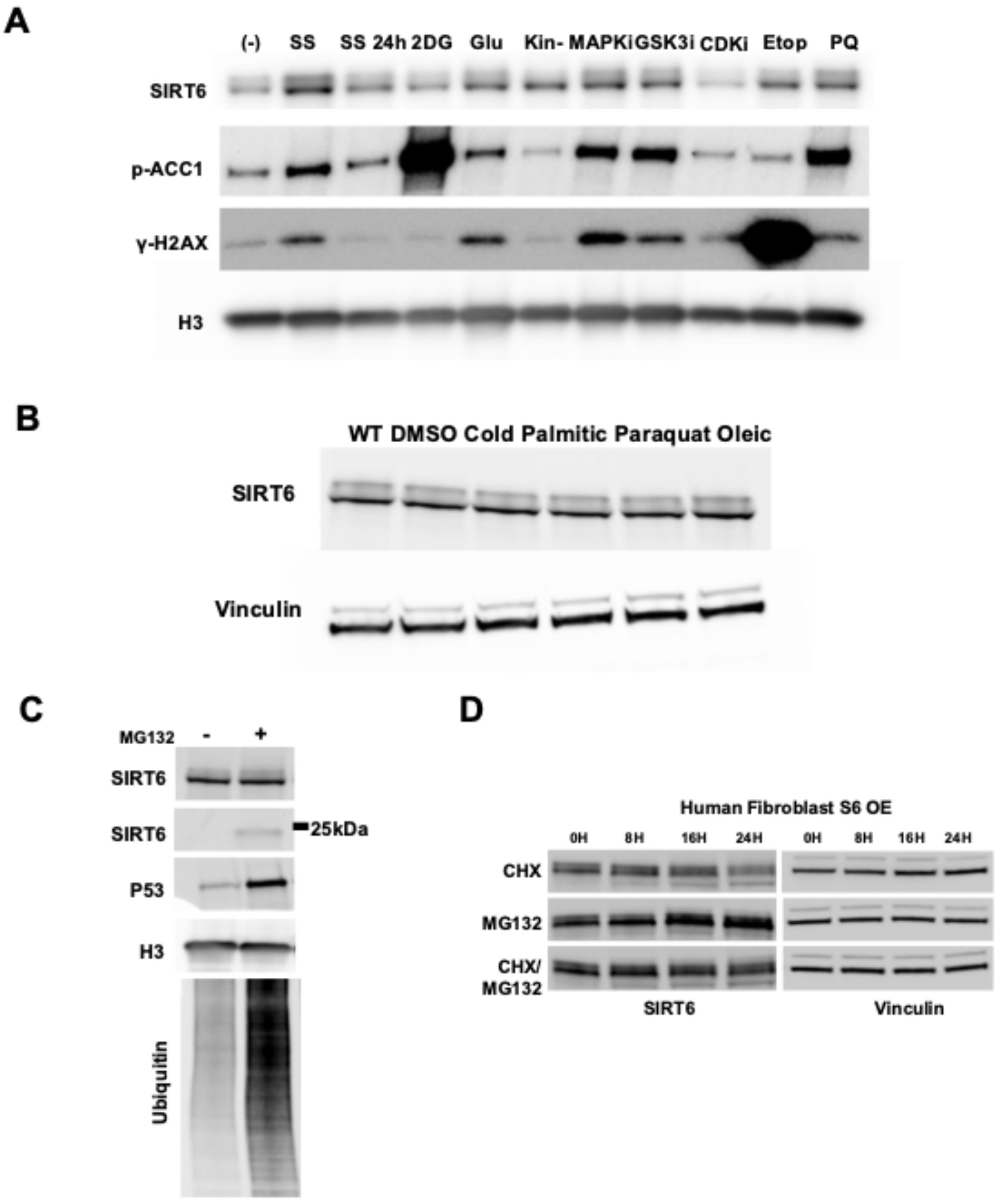
Multiple CMGC kinases progressively phosphorylate SIRT6. **A)** Western blots of fibroblast cell cultures after various treatments. Treatments were 24h unless otherwise indicated. SS = serum starvation (time indicated), 2DG = 5mM 2-deoxyglucose, Glu = 5 g/L glucose, Kin-I = Nonspecific kinase inhibition with dorsomorphin[60, 61], MAPKi = MAPK inhibition, GSKi = GSK-3 inhibition, CDKi = CDK inhibition, Etop = etoposide, PQ = paraquat. **B)** Western blots of fibroblast cell cultures after various treatments. **C)** Western blot from human fibroblasts treated with or without 10um MG132 for 24 hours. **D)** Western blot from human S6KO fibroblasts constitutively overexpressing human SIRT6 cells treated with cycloheximide, MG132, or both for the indicated time periods.

**Figure S4.**
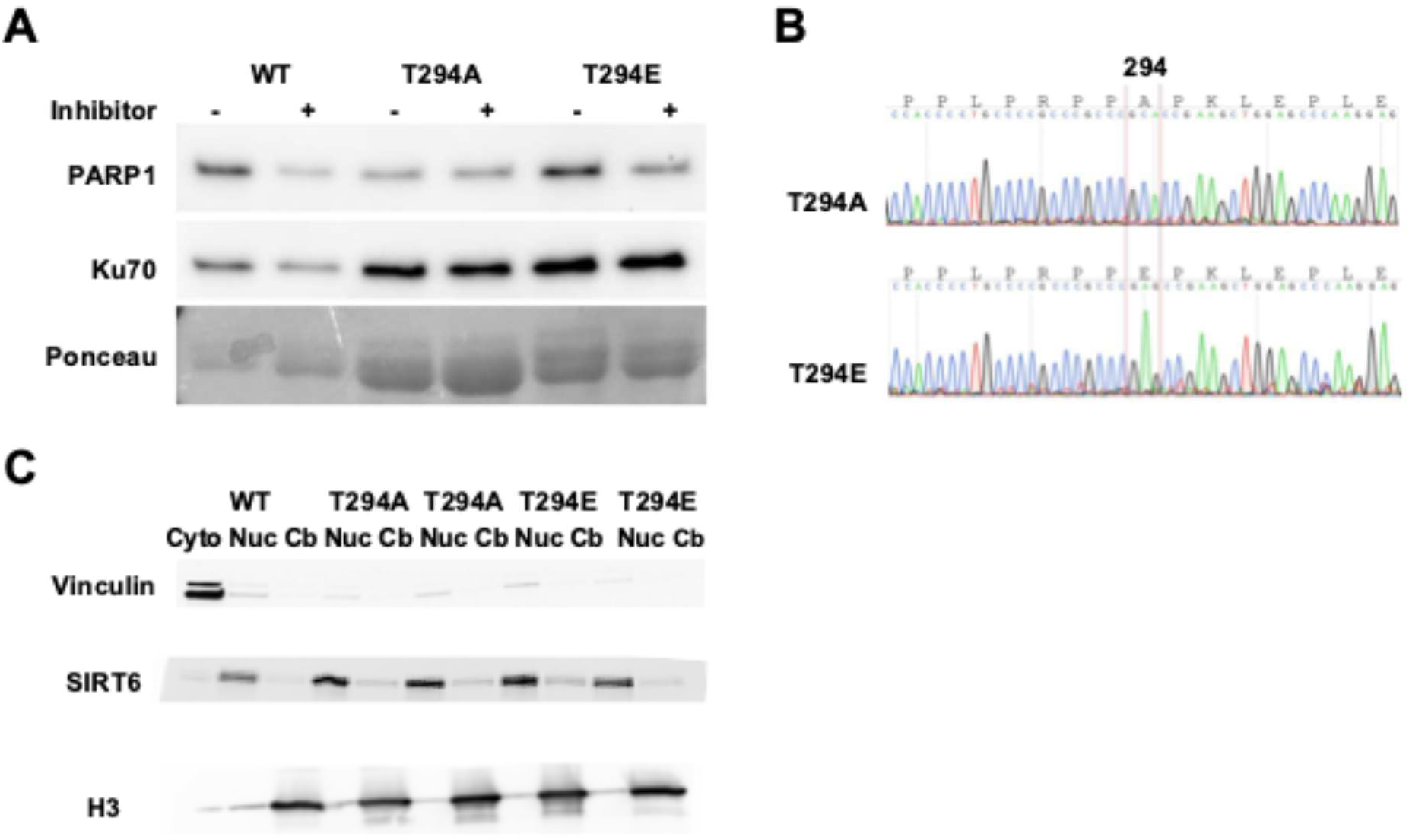
Hyperphosphorylation enhances oxidative stress resistance via PARP1. **A)** Western blots on recombinant human SIRT6 with mutation to T294 and with or without kinase inhibition during protein production. **B)** Example Sanger sequencing traces from one clone each of the T294A and T294E genotypes. In-frame codon translations are shown above the nucleotide sequence for reference. **C)** Western blots of fractionated cell lysates from human fibroblast cell lines harboring homozygous mutant endogenous SIRT6 alleles for WT, T294A “A”, or T294E “E” SIRT6. Replicate genotypes are independent clones.

**Figure S5.**
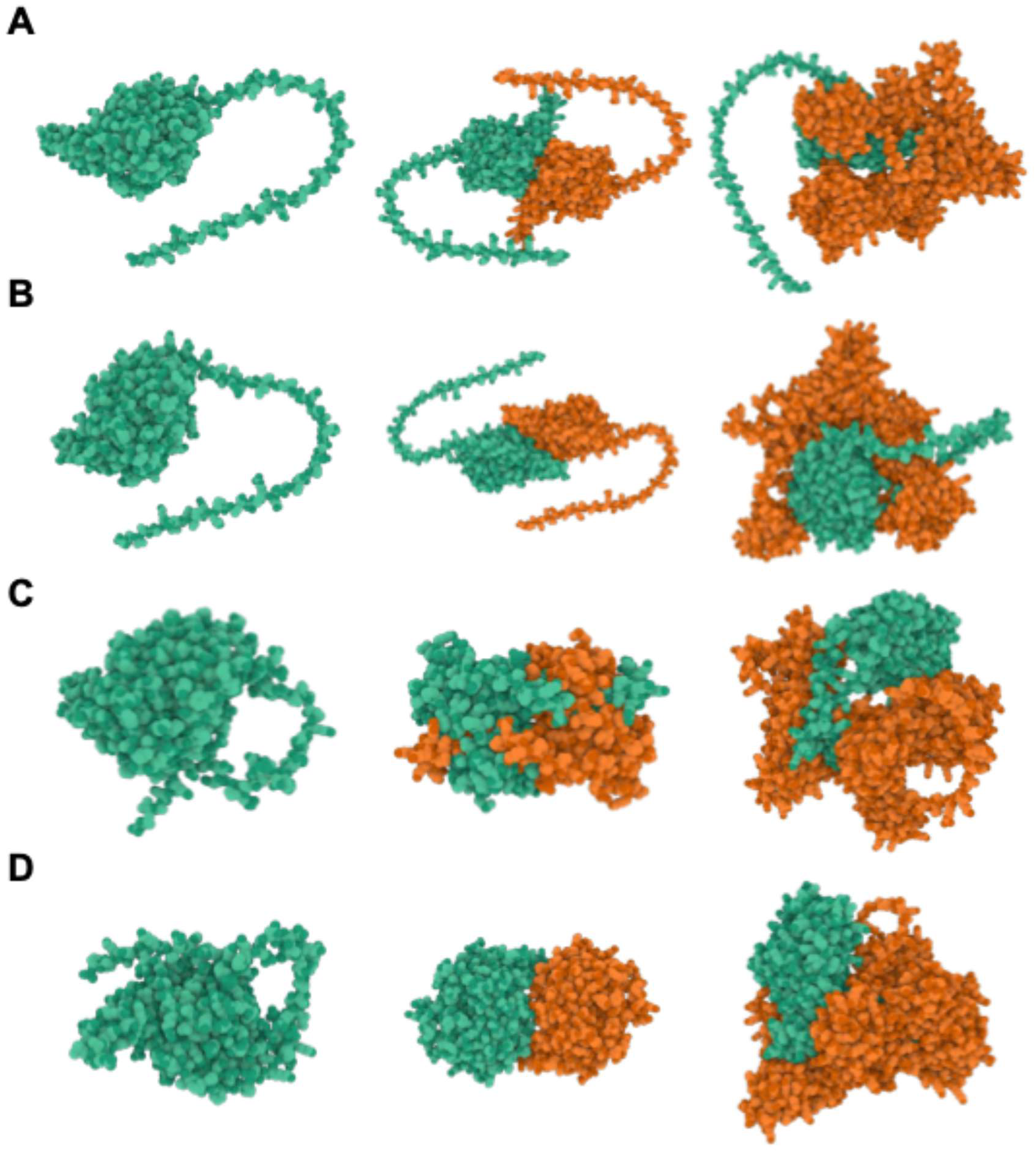
Hyperphosphorylation is predicted to enhance PARP1 interaction. **A)** AlphaFold3 structures of unmodified SIRT6 as monomer, dimer, or heterodimer with PARP1. **B)** AlphaFold3 structures of pS10 SIRT6 as monomer, dimer, or heterodimer with PARP1. **C)** AlphaFold3 structures of SIRT6 with seven C-terminal phosphorylations (T294, S303, T305, S326, S330, T337, S338) as monomer, dimer, or heterodimer with PARP1. **D)** AlphaFold3 structures of SIRT6 with pS10 and all seven C-terminal phosphorylations as monomer, dimer, or heterodimer with PARP1.

